# Acetylation of cytidine residues boosts HIV-1 gene expression by increasing viral RNA stability

**DOI:** 10.1101/2020.01.31.925578

**Authors:** Kevin Tsai, Ananda Ayyappan Jaguva Vasudevan, Cecilia Martinez Campos, Ann Emery, Ronald Swanstrom, Bryan R. Cullen

## Abstract

Covalent modifications added to individual nucleotides on mRNAs, called epitranscriptomic modifications, have recently emerged as key regulators of both cellular and viral mRNA function^1,2^ and RNA methylation has now been shown to enhance the replication of human immunodeficiency virus 1 (HIV-1) and several other viruses^3–11^. Recently, acetylation of the N^4^ position of cytidine (ac4C) was reported to boost cellular mRNA function by increasing mRNA translation and stability^12^. We therefore hypothesized that ac4C and N-acetyltransferase 10 (NAT10), the cellular enzyme that adds ac4C to RNAs, might also have been subverted by HIV-1 to increase viral gene expression. We now confirm that HIV-1 transcripts are indeed modified by addition of ac4C at multiple discreet sites and demonstrate that silent mutagenesis of a subset of these ac4C addition sites inhibits HIV-1 gene expression in *cis*. Moreover, reduced expression of NAT10, and the concomitant decrease in the level of ac4C on viral RNAs, inhibits HIV-1 replication by reducing HIV-1 RNA stability. Interestingly Remodelin, a previously reported inhibitor of NAT10 function^13,14^, also inhibits HIV-1 replication without affecting cell viability, thus raising the possibility that the addition of ac4C to viral mRNAs might emerge as a novel cellular target for antiviral drug development.

## Introduction

Previously, we and others have reported the detection and mapping of several epitranscriptomic modifications on HIV-1 transcripts^3,4,10,15^. These modifications include methylation of the N^6^ position of adenosine (m^6^A), of the C^5^ position of cytidine (m^5^C) and of the ribose moiety of all four ribonucleotides (2’O-methylation, collectively N_m_). All three of these epitranscriptomic modifications have now been shown to boost HIV-1 replication in *cis*. Specifically, m^6^A has been reported to increase viral RNA expression^3,4^, while m^5^C boosts viral mRNA translation^10^ and N_m_ residues increase HIV-1 replication by inhibiting activation of the cellular innate immune factor MDA5 by viral RNAs^15^. While m^6^A, m^5^C and N_m_ have therefore all been shown to increase HIV-1 gene expression, several other epitranscriptomic mRNA modifications remain unexamined. Of particular interest is the novel mRNA modification ac4C, which was recently reported to enhance cellular mRNA translation and stability^12^ and which has previously been reported to represent ~0.5% of all nucleotides (~2% of “C” residues) on the HIV-1 genomic RNA (gRNA), which would equate to ~8 ac4C residues per gRNA^16^. We therefore hypothesized that addition of ac4C residues to HIV-1 transcripts might also serve to boost HIV-1 gene expression and replication.

## Results & Discussion

We have previously used photo-assisted (PA) crosslinking of modification-specific antibodies to 4-thiouridine (4SU)-labelled RNA, followed by RNase footprinting of antibody-bound RNA and deep sequencing of bound RNA fragments, to map both m^6^A (PA-m^6^A-seq) and m^5^C (PA-m^5^C-seq) residues on the gRNAs and mRNAs encoded by HIV-1 and other viruses^3,5,6,10^. As an ac4C-specific antibody recently became available, we asked if a similar approach (PA-ac4C-seq) might also allow us to map ac4C modification sites on HIV-1 transcripts^17^. In Fig. 1A, we report the PA-ac4C-seq analysis of HIV-1 gRNA, isolated from HIV-1 virions generated in the CEM-SS or SupT1 T cell line, or on intracellular viral RNAs, isolated from CEM-SS cells. Despite some degree of variability in signal intensity, we were able to identify ~11 conserved sites of ac4C addition on HIV-1 transcripts that were detected across these three replicates and on additional replicates performed in HIV-1 infected CEM cells (Fig. S1), but not in mock infected cells (Note that the relative weakness of the ac4C sites detected on intracellular RNAs in the *gag, pol* and *env* regions is expected as these regions are removed by splicing in many intracellular viral RNAs). Analysis of the location of ac4C sites across the host cell transcriptome confirmed the previous report that the majority of the mapped ac4C residues are located in coding sequences (CDS)^12^, though a substantial number were also located in 3’ untranslated regions (UTR) (Fig. S1B). Of interest, analysis of all mapped ac4C sites identified a C and U-rich consensus sequence, with a central “UCU” motif, in both uninfected and HIV-1 infected CEM cells (Fig. S1C).

**Fig 1.**
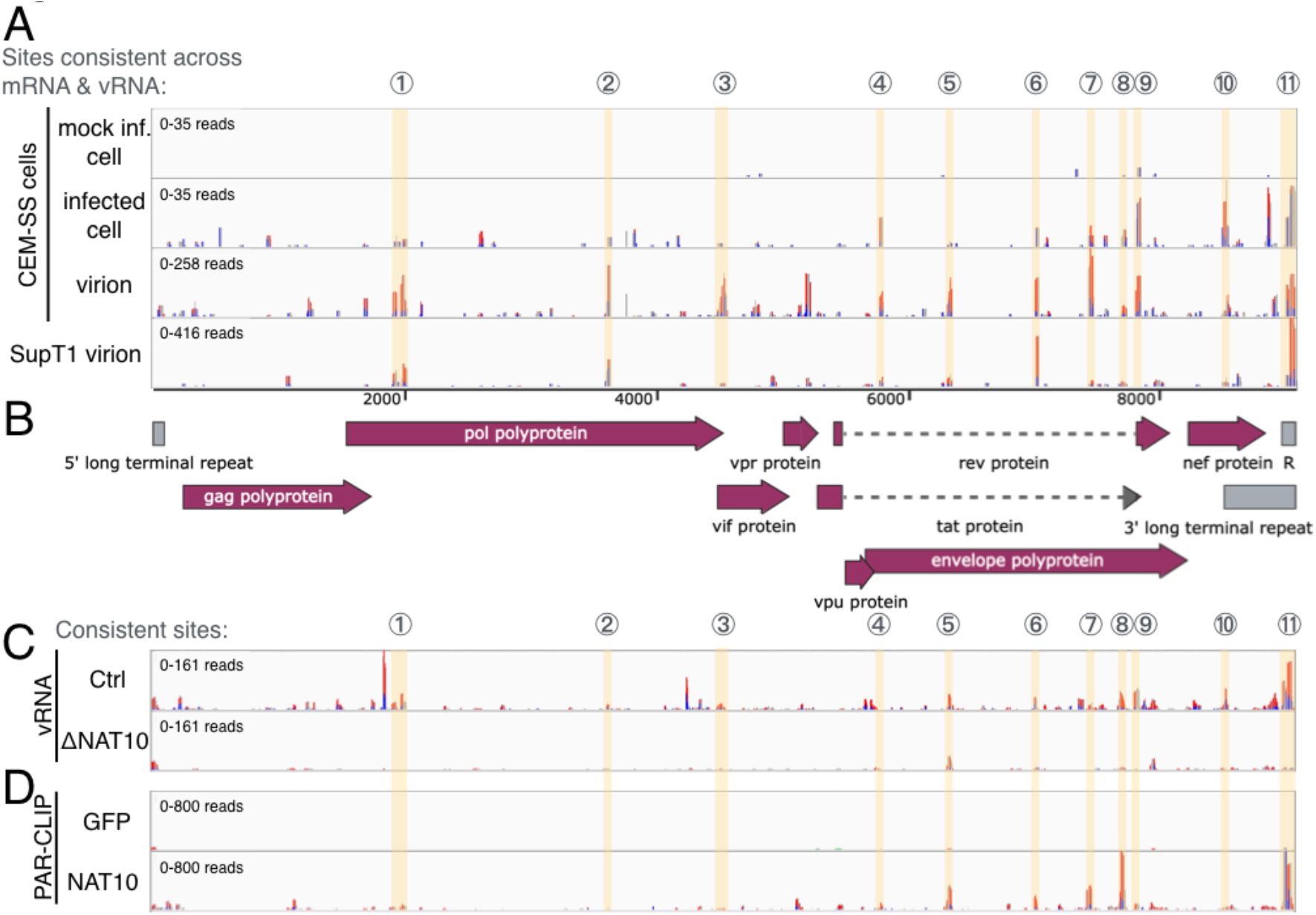
NAT10-dependent ac4C is deposited at multiple locations on HIV-1 mRNAs and virion genomic RNA. (A) Ac4C sites were mapped by PA-ac4C-seq on the poly(A)^+^ fraction (mRNA) of mock or HIV-1 infected CEM-SS T cells, along with virion particle RNA produced from HIV-1 infected Sup T1, and CEM-SS cells. (B) Schematic of the HIV-1 genome organization drawn to scale (C) PA-ac4C-seq was performed on HIV-1 virion RNA produced in control or ΔNAT10 CEM T cells. NAT10 knock-down is validated in Fig. 3A. (D) PAR-CLIP was performed on CEM T cells stably expressing FLAG-NAT10 or FLAG-GFP to identify NAT10 binding sites on HIV-1 RNA. Sequence reads were aligned to the HIV-1 NL4-3 genome. Consistent (across mRNA and virion) ac4C sites highlighted in yellow and numbered above. 4SU-based CLIP methods result in T>C conversions where protein is cross-linked to 4SU residues, here shown as red-blue bars.

If the data reported in Fig. 1A indeed map authentic ac4C addition sites, then loss of the “writer” acetyltransferase that deposits these marks should result in the loss of detectable ac4C residues on viral RNAs. Mammals express a single RNA acetyltransferase capable of acetylating RNA, N-acetyltransferase 10 (NAT10), and NAT10 has indeed been reported to add ac4C to mRNAs and non-coding RNAs in human cells^12,18,19^. However, NAT10 appears to be essential in human cells and previous efforts to knock down NAT10 expression using gene editing by CRISPR/Cas have therefore focused on an exon, exon 5, that is found in the majority of, but not all, mRNAs encoding NAT10^12^. We repeated this strategy in the T cell line CEM and generated 3 clonal cell lines in which all three copies of the NAT10 gene were edited in exon 5 (Figs. S2A, B and C), resulting in a large decrease in NAT10 expression (see below). Of note, these clonal cell lines, referred to as ΔNAT10 #3, #7 and #9, did not show any decrease in growth rate (Fig. S2D). However, reduced NAT10 expression did result in the expected strong decline in the level of ac4C residues present on viral transcripts as measured by PA-ac4C-seq, thus validating these mapped ac4C sites as authentic (Fig. 1C).

If NAT10 is indeed the writer that deposits ac4C on cellular and viral RNAs, then we reasoned it might be possible to detect NAT10 binding to these sites using the photo-assisted cross linking and immunoprecipitation (PAR-CLIP) technique, as previously described^3,20^. We therefore used lentiviral expression vectors encoding FLAG-tagged wild type (WT) NAT10, or FLAG-tagged green fluorescent protein (GFP), to stably express these proteins in CEM cells. As shown in Fig. 1D, we indeed detected FLAG-NAT10, but not FLAG-GFP, binding sites on viral transcripts and these were coincident with the mapped ac4C sites, as expected. Moreover, the mapped NAT10 binding sites conformed to the same U and C-rich sequence consensus, with a central “UCU” motif, that we had identified using PA-ac4C-seq (Fig. S1C).

If ac4C residues indeed facilitate some aspect of HIV-1 gene expression then the reduced expression of NAT10, and concomitant reduction in ac4C addition to mRNAs, seen in the ΔNAT10 CEM cells should result in reduced HIV-1 replication. We analyzed the level of HIV-1 Gag and NAT10 protein expression in WT and ΔNAT10 CEM cells 3 days after infection using WT HIV-1 isolate NL4-3. As may be observed in Fig. 2A, both NAT10 and the viral Gag proteins are expressed at a much lower level in the ΔNAT10 CEM cells, though the GAPDH loading control was unaffected. Similarly, when we infected WT and ΔNAT10 CEM cells with a previously described replication competent HIV-1 that encodes the Nano luciferase (*NLuc*) gene in place of the dispensable *nef* gene^21^, we observed a strong reduction in the level of NLuc protein expression (Fig. 2B) and in the level of viral RNA expression, measured by qRT-PCR using an LTR-specific probe (Fig. 2C), in the latter. Thus, reduced NAT10 expression indeed results in a lower level of HIV-1 replication.

**Fig 2.**
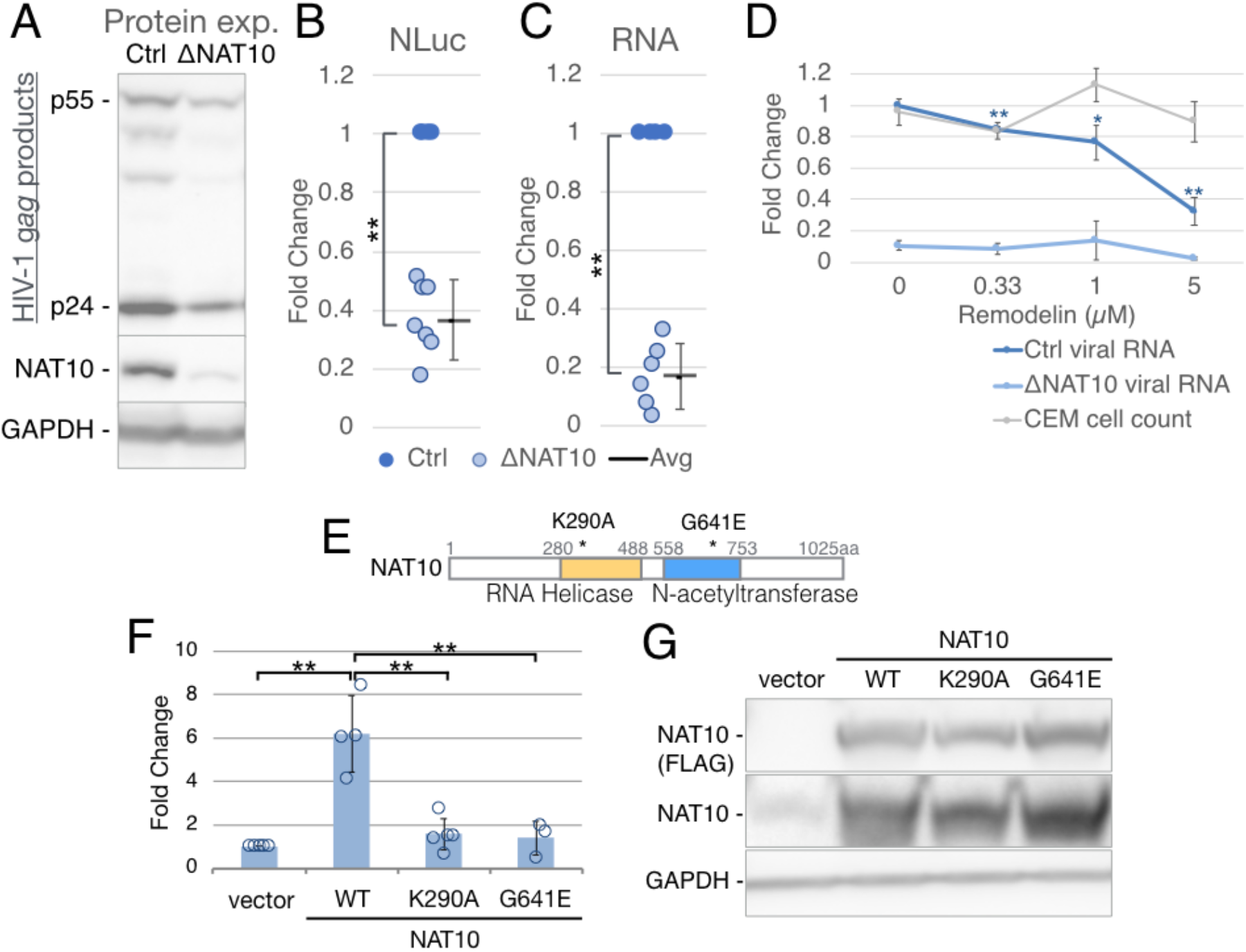
NAT10 enhances the rate of HIV-1 spread in culture. (A-C) CEM cells in which the *NAT10* gene had been edited using CRISPR/Cas (ΔNAT10), along with control CEM cells expressing Cas9 and a non-targeting guide RNA (Ctrl), were assayed for NAT10-associated viral replication phenotypes. (A) HIV-1 replication levels in a spreading infection, at 3 dpi with WT NL4-3, were analyzed by Western blot for the HIV-1 capsid protein p24 (B) WT and ΔNAT10 cells were infected with the NL4-3NLuc reporter virus and NLuc activity determined at 3 dpi (C) HIV-1 RNA levels in the samples shown in panel B were determined using qRT-PCR. Three different ΔNAT10 single cell clones were used in panels B & C, with Ctrl cells set at 1. n=4 to 7, error bars=SD. ** 2-tailed T-test, p<0.01. (D) CEM, Ctrl, and ΔNAT10 cells were treated with the NAT10-inhibitor Remodelin. Infected CEM cells were counted at 1 dpi to determine Remodelin toxicity (shown in gray), infected Ctrl (dark blue) and ΔNAT10 (light blue) cells were harvested at 2 dpi and viral RNA levels assayed by qRT-PCR. n=3, error bars=SD. 2-tailed T-test on Ctrl cells for each condition compared to the 0 μM level, **p<0.01, *p<0.05. (E) Schematic of NAT10 functional domains and point mutations. (F) 293T cells were transfected with empty vector, WT NAT10 (WT), K290A or G641E mutant expression vectors. 3 days later, transfected cells were infected with the NL4-3NLuc reporter virus, and NLuc assayed 2 dpi. n=3 to 6, error bars=SD. ** 2-tailed T-test, p<0.01. (G) Western blot showing NAT10 over-expression, NAT10 probed with both FLAG and NAT10 antibodies and GAPDH probed as a loading control.

While these data address how lower NAT10 expression affects HIV-1 replication, it is also possible to inhibit NAT10 function in WT cells using a drug, called Remodelin, that has been reported to inhibit NAT10 function at concentrations that are non-toxic in culture or in mice^13,14^. Indeed, we observed that Remodelin reduced HIV-1 replication in WT CEM cells by up to 70%, but had little effect on HIV-1 replication in the ΔNAT10 CEM cells, at concentrations that did not reduce CEM cell growth (Fig. 2D), thus further validating NAT10 as an HIV-1 co-factor.

As reduced NAT10 expression or function led to diminished HIV-1 replication (Figs. 2A-D), we reasoned that NAT10 activity might be rate limiting for HIV-1 replication in CEM cells. We therefore generated CEM cells stably overexpressing WT NAT10, or mutant forms of NAT10 lacking a functional RNA helicase domain (K290A) or acetyltransferase domain (G641E) (Fig. 2E), due to mutagenesis of residues previously shown to be required for RNA acetyltransferase function^14,19^. All three proteins were expressed at similar levels and at levels that were much higher than endogenous NAT10, as determined by Western blot (Fig. 2G). Importantly, we detected 4-8x higher levels of HIV-1 replication in the CEM cells overexpressing WT NAT10, but not with either NAT10 mutant, when compared to the parental CEM cells and this difference was highly significant (Fig. 2F).

While the data presented in Fig. 2 demonstrate that NAT10, and the ac4C modification, promote some aspect(s) of the HIV-1 replication cycle, they do not identify which step(s) are affected. To address this issue, we performed a single cycle HIV-1 replication assay, using WT NL4-3, in WT or ΔNAT10 cells and measured the efficiency of several different steps in the HIV-1 replication cycle. Initially, we measured the level of viral Gag expression, which was found to be reduced by ~70%. Measurement of total viral RNA expression, by qRT-PCR using an LTR-specific probe, also revealed an ~70 % reduction in the ΔNAT10 CEM cells when compared to WT, suggesting an effect primarily at the RNA level (Fig. 3A). Indeed, analysis of the level of ribosome binding by viral mRNAs^22^, an assay which had revealed a strong positive effect of the m^5^C modification on HIV-1 mRNA translation^10^, indicated that the presence or absence of ac4C had no discernable effect (Fig. 3C). Similarly, reduced ac4C addition did not affect the subcellular location of HIV-1 transcripts (Fig 3D), or their alternative splicing (Fig. S3A). Importantly, none of the steps from cell entry, reverse transcription to proviral integration were affected by loss of ac4C, as no difference was found in the total level of HIV-1 DNA (Fig. S3B). However, the reduced expression of NAT10, and the concomitant loss of ac4C on viral transcripts, did result in a highly significant reduction in the stability of HIV-1 transcripts measured either by pulse-chase, using 4SU incorporation into RNA (Fig. 3E)^23,24^, or by use of the transcription inhibitor actinomycin D (Fig. S3C).

**Fig 3.**
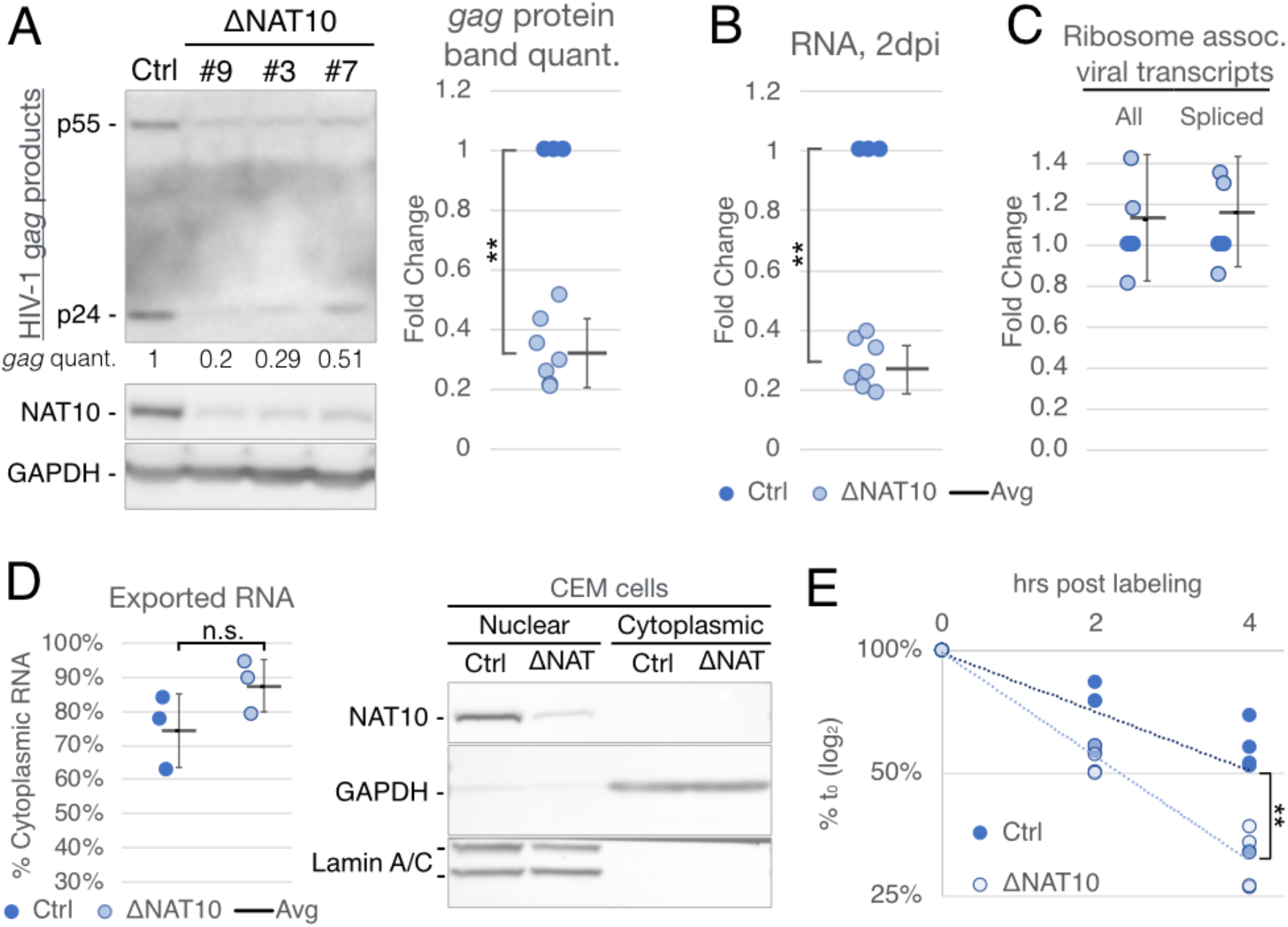
NAT10 depletion reduces HIV-1 RNA levels by destabilizing viral transcripts. Ctrl (dark blue) and ΔNAT10 (light blue) cells were infected with HIV-1, treated with the reverse transcriptase (RT) inhibitor Nevirapine (NVP) at 15-16 hpi and harvested at ~48 hpi for the following single cycle infection assays. (A) Viral Gag levels assayed on Ctrl and 3 single cell clones (#9, #3, #7) of ΔNAT10 cells by Western blot. HIV-1 Gag band intensities (p24 plus p55) were quantified, normalized to Ctrl levels (set to 1), and are shown in the right hand panel (Ctrl n=3, ΔNAT10 n=7). (B) Aliquots of the samples visualized in panel A were assayed for viral RNA levels by qRT-PCR. (C) Ribosome-associated viral RNAs were extracted and assayed by qRT-PCR using primer sets targeting the LTR U3 region (All) and across the D1 to A1 viral splice junction (Spliced), n=3. (D) Subcellular fractionation of infected Ctrl & ΔNAT10 cells. Viral RNA in each fraction was quantified by qRT-PCR and is shown as the percentage of cytoplasmic RNA over total (cytoplasmic + nuclear) RNA, n=3. Fractionation validated by Western blot in the right panel, with Lamin A/C as the nuclear marker and GAPDH as the cytosolic marker. Statistical analyses shown in (A-D) used the two-tailed Student’s T test, error bars=SD, **p<0.01. (E) Viral RNA stability tested by 4SU-metabolic labeling, followed by isolation of 4SU^+^ nascent RNA at 0, 2 and 4hrs post 4SU-labeling, n=5. Slopes of regression lines compared by ANCOVA, **p=0.0008.

The data presented in Fig 3 argue that, in the case of HIV-1 RNAs, ac4C acts to increase viral gene expression primarily by enhancing viral RNA stability. These data contrast with the previous work proposing that ac4C increases cellular mRNA gene expression by not only increasing mRNA stability but also translation by increasing the CDS decoding efficiency^12^. If this is indeed the primary mechanism of action of ac4C, then only ac4C sites present in the CDS should affect mRNA function in *cis*. To address this question, we introduced as many silent C to U mutations as possible into conserved ac4C peaks 4 through 8 in the viral *env* gene region (Fig. 4A and Figs. S4A and B). We then transfected WT 293T cells, which lack CD4 and therefore will not support a spreading infection, and measured HIV-1 Gag protein expression. As shown in Figs. 4B and 4C, the ac4C site mutations introduced into the *env* gene, which would be present exclusively in the 3’ UTR of the viral *gag* mRNA (Fig. 1B), nevertheless reduced Gag protein expression in *cis*, both in the producer cells (Fig. 4B) and in the supernatant media (Fig. 4C). The observed reduction of ~60% was not only highly significant but also only slightly less than seen in the ΔNAT10 CEM cells (Fig. 3A).

**Fig 4.**
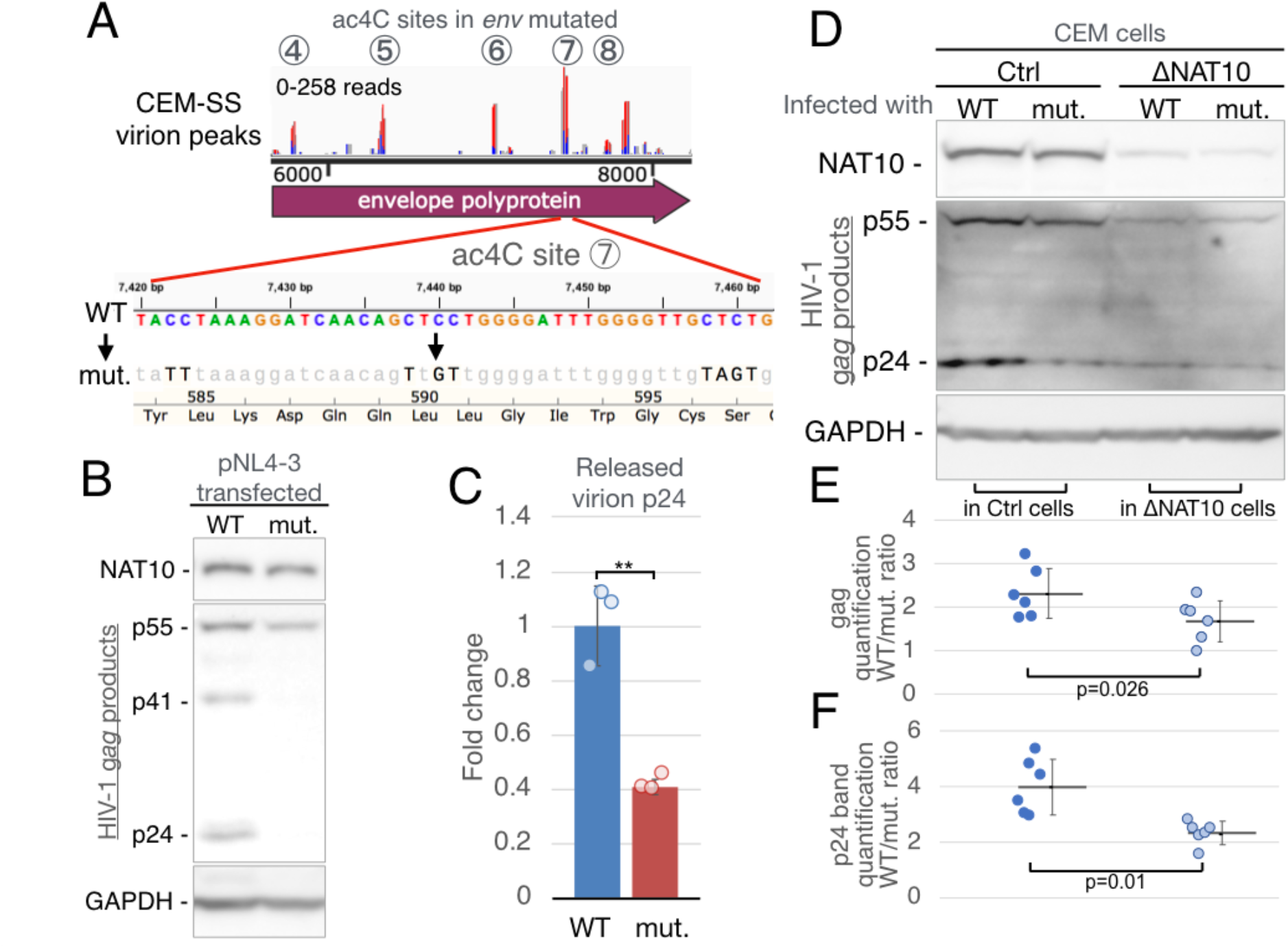
Silent mutagenesis of ac4C sites in env diminishes viral Gag expression. (A) ac4C sites #4-8 in the HIV-1 env CDS were mutated to remove as many ac4C sites as possible without changing the encoded amino acid. Example of silent mutations in ac4C site #7 in the lower panel. (B) 293T cells were transfected with WT pNL4-3 HIV-1 (WT) or the mutant viral plasmid (mut.) with ac4C sites in the env gene mutated and Gag expression determined by Western blot. (C) Virions released into the supernatant media of 293T cells transfected with WT or mut HIV-1 expression plasmids were quantified by p24 ELISA. n=4, **p=0.002 (D) WT or ac4C mut. virus were normalized using the p24 levels determined panel C, and used to infect Ctrl & ΔNAT10 CEM cells. Viral Gag expression from single round (NVP-treated) infections were assayed at 48 hpi by Western blot. (E) The gag protein bands (p55+p24) from 6 repeats of panel D were quantified and the WT/mut ratio from Ctrl & ΔNAT10 CEMs plotted. (F) Similar to panel F, WT/mut. ratios of the p24 bands only. Significance determined using paired two-tailed Student’s T test, n=6, error bars=SD.

It could be argued that the inhibition of Gag expression seen with the *env* gene ac4C mutant (Figs 4B and C) was not due to loss of ac4C residues but rather due to disruption of some other sequence element that is functionally significant. To test this idea, we collected the WT and mutant virions released by the transfected 293T cells, normalized the p24 level based on Fig. 4C and then infected WT and ΔNAT10 CEM cells. We then measured the level of Gag expression after a single round of replication in these cells by Western. A representative experiment is shown in Fig. 4D, while a compilation of data measuring total Gag protein expression (Fig. 4E) or exclusively p24 Gag expression (Fig. 4G) are also presented. We noted a bigger effect on p24 Gag expression than on total Gag protein expression and therefore present both data sets. As may be observed, the *env* ac4C site mutations reduced total Gag protein expression by 2.3x, and p24 Gag expression by 4.0x, in the WT CEM cells. In contrast, these same mutations reduced total Gag protein expression by 1.7x, and p24 expression by 2.1x, in the ΔNAT10 CEM cells, and these differences are statistically significant. Therefore, these data demonstrate that the mutagenesis of mapped *env* ac4C sites indeed results a stronger inhibitory phenotype in CEM cells expressing WT levels of NAT10 than in the ΔNAT10 CEM cells that express reduced levels of NAT10, as would be predicted if they indeed act via the same mechanism.

Previously, several groups have reported that the epitranscriptomic addition of m^6^A to viral transcripts can significantly enhance the replication of a range of different viruses, including HIV-1, influenza A virus, SV40, enterovirus 71, respiratory syncytial virus and Kaposi’s sarcoma-associated herpesvirus^3–9,11^. Less is known about other epitranscriptomic viral RNA modifications, though both m^5^C and N_m_ residues have been detected on HIV-1 transcripts at levels that are substantially higher than seen on cellular mRNAs and both m^5^C and N_m_ have been reported to enhance HIV-1 replication in culture^10,15^. Interestingly, the proposed mechanisms used by these distinct epitranscriptomic modifications appear distinct in that m^6^A has been proposed to increase viral mRNA expression levels^3,4^ while m^5^C acts primarily by boosting viral mRNA translation^10^. Finally N_m_ has been reported to increase HIV-1 replication indirectly by preventing the activation of the host innate immune factor MDA5 by viral transcripts^15^. Here, we extend this previous work by looking at a novel epitranscriptomic modification, ac4C, that has been proposed to boost cellular mRNA translation and stability^12^ and that has also been detected on purified HIV-1 genomic RNA^16^. We have mapped the ac4C residues present on HIV-1 RNAs to ~11 distinct sites and show that these are, as expected, deposited by the host acetyltransferase NAT10, as inhibition of NAT10 expression results in a loss of ac4C from viral RNAs (Fig.1). Importantly, the loss of ac4C modifications from viral transcripts results in reduced viral gene expression and replication whether caused by a reduction in NAT10 expression due to gene editing (Figs. 2A and B and Fig. 3), inhibition of NAT10 function using the drug Remodelin (Fig. 2D) or by mutagenesis of mapped ac4C sites (Fig. 4). However, in the case of HIV-1 transcripts, the positive effect of ac4C modifications appears to be due entirely to stabilization of viral transcripts (Figs. 3E and S3C). In contrast, Arango et al.^12^ reported that ac4C residues in cellular CDS not only increased mRNA stability but also increased mRNA translation, by increasing decoding efficiency. We did not observe any increase in ribosome recruitment to HIV-1 mRNAs (Fig. 3C), a result that contrasts with what we observed for m^5^C, which in our hands clearly enhanced HIV-1 mRNA translation^10^. Moreover, we observed that mutagenesis of ac4C sites in the viral *env* gene, which forms part of the 3’ UTR of viral mRNAs encoding Gag, nevertheless resulted in a marked drop in Gag protein expression (Figs. 4B and C). It remains unclear whether this indicates that ac4C acts differently on HIV-1 and cellular mRNAs or whether HIV-1 mRNAs are already maximally optimized for ribosome decoding and therefore this parameter cannot be further enhanced by ac4C. Regardless, these data do clearly demonstrate that NAT10 adds ac4C to HIV-1 transcripts at multiple discrete locations and identify ac4C as the fourth epitranscriptomic modification to enhance viral replication in *cis*.

## Methods

### Cell lines

CEM, CEM-SS and SupT1 are CD4+ T cell lines that were obtained from the NIH AIDS reagent program. T cell lines were cultured in Roswell Park Memorial Institute (RPMI) 1640 medium supplemented with 10% fetal bovine serum (FBS) and 1% Antibiotic-Antimycotic (Gibco, 15240062). 293T cells were cultured in Dulbecco’s Modified Eagle’s Medium (DMEM) with 6% FBS and 1% Antibiotic-Antimycotic. ΔNAT10 CEM cells were produced by transducing CEM cells with a lentiviral vector, LentiCRISPRv2^25^, encoding Cas9 and single guide RNAs (sgRNAs) 5’-TGAGTTCATGGTCCGTAGG-3’ (as previously published^12^) or 5’-GGCTAGTGGTCATCCTCCTA-3’ (GeCKOv2, guide number HGLibA_31166). Two days post-transduction, cells were subjected to 1-2 weeks of selection in 1 μg/ml puromycin, then single cell cloned by limiting dilution. Control CEM cells were produced by transduction with a lentiviral vector expressing a non-targeting sgRNA specific for GFP (5’-GTAGGTCAGGGTGGTCACGA-3’), then puromycin selected and single cell cloned. To validate CRISPR mutations, genomic DNA from knockdown cell lines was extracted using the Zymo Quick-DNA Miniprep Plus kit (#11-397). The genomic region flanking the Cas9 target site from each ΔNAT10 cell line was PCR amplified and cloned into the XbaI/SalI sites of pGEM-3zf+ vector (Promega). 10+ bacterial cell clones of pGEM-genomic-region-plasmid from each CRISPR-knockdown cell clone were isolated for Sanger sequencing. CEM cells constitutively expressing FLAG-NAT10 or FLAG-GFP were produced using lentiviral expression vectors pLEX-FLAG-NAT10 (as described below) or pLEX-FLAG-GFP (previously described^3^), following the same transduction and single cell selection process as above.

### Antibodies

Antibodies used in this study include: Anti-ac4C, a generous gift from Dr. Shalini Oberdoerffer (NIH, NCI)^17^, a later batch was purchased from Abcam ab252215. NAT10, Proteintech 13365-1-AP. FLAG antibody clone M2, Sigma F1804. GAPDH, Proteintech 60004-1-Ig. β-Actin, Proteintech 66009-1-Ig. Lamin A/C clone E-1, Santa Cruz, sc-376248. Anti-Mouse HRP, Sigma A9044. Anti-Rabbit HRP, Sigma A6154. HIV-1 p24 Gag Monoclonal (#24-3) from Dr. Michael H. Malim^26^.

### Viruses

Recombinant virus clones used include the laboratory strain NL4-3 (NIH AIDS Reagent, #114)^27^ and the nano-luciferase reporter virus NL4-3-NLuc^21^. A mutant NL4-3 virus with most ac4C sites in *env* silently mutated was cloned by replacing the SalI-NheI and NheI-BamHI segments of *env* with gBlocks (IDT) designed to mutate any C in identified ac4C peaks that could be mutated without changing the encoded amino acid. All viruses were produced by transfecting the viral expression plasmid into 293T cells using polyethylenimine (PEI, 2 μg plasmid for a 6 well plate, 10 μg for a 10cm plate, 20μg for a 20cm plate, PEI used at 2.5x the μg amount of plasmid DNA). 24 hours post-transfection, the media were replaced with fresh media. The supernatant media were collected at 72 hours post-infection (hpi), passed through a 0.45μm filter, then overlaid onto target cells.

### PA-ac4C-seq

Harvest of cellular and virion RNA for modification mapping was performed as previously reported^10^. CEM or SupT1 cells were resuspended in filtered supernatant media from NL4-3-transfected 293T cells. Mock-infected cells were resuspended in filtered media from non-transfected 293T cells. The media were replaced with fresh media at 24 hpi to reduce carry over of 293T-produced virions. Cells were pulsed with 100μM 4SU at 48 hpi and incubated an additional 24 hours. At 72 hpi, cells were collected for extraction of total RNA using Trizol (Invitrogen), followed by mRNA enrichment using the Poly(A)Purist MAG Kit (Ambion). For virion RNA, the supernatant of infected cells was concentrated through Centricon Plus-70 centrifugal filters (100,000 NMWL membrane, Millipore), then the virions were pelleted by ultracentrifugation through a 20% sucrose cushion at 38,000 rpm for 90 min. The resulting virus pellet was lysed in Trizol for RNA extraction. Cellular poly(A)+ RNA and virion RNA were then subjected to ac4C site recovery following the PA-m^6^A-seq protocol^6,10,28^. with two modifications: an ac4C-specific antibody was used, and the incubation of antibody with RNA was overnight.

### PAR-CLIP

Single cell cloned CEM cells expressing FLAG-GFP or FLAG-NAT10 were used. Ten 15 cm plates of 293T cells were seeded to package virus for each (GFP+ or NAT10+) infection. Four days post pNL4-3 transfection, the virus-containing supernatant media were harvested and filtered through a 0.45μM filter. 300 million FLAG-GFP or FLAG-NAT10 expressing CEM cells were resuspended in the filtered media. At 48 hp, infected cells were pelleted and resuspended in 350 ml of fresh RPMI supplemented with 100μM 4SU. At 72 hpi, cells were collected, washed twice in PBS, then irradiated with 2500×100 μJ/cm^2^ of 365 nm UV. The above procedure was repeated twice to obtain sufficient biomass to perform PAR-CLIP, as previously described^3,20^, using an anti-FLAG antibody.

### Illumina sequencing & bioinformatic data analysis

RNA recovered from the PA-ac4C-seq and PAR-CLIP procedures were used for cDNA library preparation using the NEBNext Small RNA Library Prep Set for Illumina (NEB E7330S), then sequenced using Illumina NextSeq 500, or NovaSeq 6000 sequencers at the Duke Center for Genomic and Computational Biology (GCB) Sequencing and Genomic Technologies Shared Resource. Sequencing data analysis was done as previously described^6,10^. Sequencing reads >15 nt with fastq quality score >33 were first aligned to the human genome (hg19) using Bowtie^29^. The human-non-aligning reads were then aligned (allowing up to 1 mismatch) to the HIV-1 NL4-3 sequence with a single copy of the long terminal repeat (LTR, U5 on the 5’ end, and U3-R on the 3’ end), essentially 551-9626 nt of GenBank AF324493.2. As UV-crosslinked 4SU results in characteristic T>C conversions, an in-house Perl script was used to discard alignments devoid of T>C mutations. After file format conversions using SAMtools^30^, data was visualized using IGV^31^. For meta-gene analysis and motif analysis, the human-aligned PA-ac4C-seq reads were subjected to peak calling using MACS2^32^ (parameters -- nomodel --tsize=50 --extsize 32 --shift 0 --keep-dup all-g hs). Peak calling on the NAT10 PAR-CLIP data was done using PARalyzer v1.1^33^ (with the parameters:

BANDWIDTH=3 CONVERSION=T>C
MINIMUM_READ_COUNT_PER_GROUP=10
MINIMUM_READ_COUNT_PER_CLUSTER=3
MINIMUM_READ_COUNT_FOR_KDE=5 MINIMUM_CLUSTER_SIZE=15
MINIMUM_CONVERSION_LOCATIONS_FOR_CLUSTER=2
MINIMUM_CONVERSION_COUNT_FOR_CLUSTER=2
MINIMUM_READ_COUNT_FOR_CLUSTER_INCLUSION=2
MINIMUM_READ_LENGTH=10
MAXIMUM_NUMBER_OF_NON_CONVERSION_MISMATCHES=1
EXTEND_BY_READ).

Motif analysis was then performed using MEME following a published m^6^A-seq pipeline^34^. Metagene analysis was performed using metaPlotR^35^.

### Expression plasmid construction

A NAT10 cDNA was cloned by PCR from CEM-SS cDNA, digested and ligated into the NotI and EcoRI sites of pK-FLAG-VP1^6^, placing NAT10 3’ to a 2xFLAG tag and replacing SV40 VP1. The NAT10 cDNA sequence was confirmed as wild type by Sanger DNA sequencing. The K290A & G641E point mutants were introduced by recombinant PCR: two complementing PCR primers were designed to overlap the mutated region, with the point mutant sequence in the middle. A first round of PCR was done to separately amplify the 5’end-to-mutation site and the mutation site-to-3’end fragments of NAT10, yielding two fragments with a region of homology around the mutation site. Using two outer primers, the two fragments were joined and amplified into the full length NAT10 CDS containing the point mutation and then ligated into the NotI and EcoRI sites of pK-FLAG as before. The lentiviral expression construct pLEX-FLAG-NAT10 was constructed by cloning the PCR amplified FLAG-NAT10 cDNA from pK-FLAG-NAT10 into the BamHI and AgeI sites of the pLEX vector (Openbiosystems). All PCR primers used are listed in Supplemental Table 1.

### Viral infection of 293T cells

293T cells seeded in 6 well plates were transfected using PEI with 1.6 μg of pK-FLAG-NAT10 plasmids or empty pK vector, along with 250 ng of CD4 expression vector^36^, and 100 ng of firefly luciferase (FLuc) expression plasmid pcDNA3-FLuc. In parallel, 10μg of pNL43-NLuc was transfected into 293T cells in 10 cm plates. All media were changed the next day, and the NAT10/CD4/FLuc+ infected target cells split into 12 well plates two days later. NL43-NLuc virus-containing supernatant was harvested on day 3, filtered and brought up to 12mls with fresh media, and overlaid onto target cells at 1ml virus per well. At 2 or 3dpi, the supernatant media was removed from infected cells, the cells washed 3x with PBS, then lysed in passive lysis buffer (Promega, E1941). NLuc and FLuc activity was assayed using the Nano Luciferase Assay Kit and Luciferase Assay System (Promega, N1120 & E1500).

### Remodelin assays

Control or ΔNAT10 CEM cells were seeded at 0.75 million cells per well in 1ml in a 12 well plate, and treated with Remodelin (Sigma SML1112-5MG, dissolved in DMSO to 2mM) at the needed concentration. Lower concentration sets were compensated with equal volumes of DMSO. The next day, cells were overlaid with 1 ml of NL4-3 virus. Additional drug was added to compensate for the additional 1 ml volume of the virus. To compensate for potential drug decay over 3 days, 0.25x additional drug was added at 2 days post initial drug treatment. Cells were counted at 24 hpi to assess toxicity and harvested at 48 hpi for assay of viral RNA levels.

### Viral infection of wild type and ΔNAT10 CEM cells

NL4-3 virus was packaged in 293T cells in 10 cm plates, transfected with 10 μg of pNL4-3 using PEI, the media were replaced the next day with 10 ml RPMI. 3 days post-transfection, CEM cells were counted and seeded at 1 million cells per well in 0.5 ml RPMI in 12 well plates. Virus-containing supernatant media harvested from 293T cells were filtered and supplemented with fresh RPMI to a total volume of 12 ml. 1 ml of this virus was then used to overlay the 1 million cells/0.5 ml in 12 well plates. For spreading infections, cells were harvested and PBS washed at 72 hpi. For single-round infections, cells were treated with 133μM of the reverse transcriptase inhibitor Nevirapine (Sigma SML0097) at 16 hpi, then harvested at 48 hpi.

Viral RNA levels in infected cells were measured using quantitative real time-PCR (qRT-PCR). Harvested cells were washed in PBS, then the RNA extracted using Trizol (Invitrogen). Total RNA was then treated with DNase I (NEB), and reverse transcribed using the Super Script III reverse transcriptase (Invitrogen). qPCR was performed with Power SYBR Green Master Mix (ABI), with primers targeting either the U3 region of HIV-1 LTR or spanning splice donor 1 and splice acceptor 1 (D1-A1). qPCR readouts were normalized to GAPDH levels using the delta-delta Ct (ddCt) method. All PCR primers used are listed in Supplemental Table 1. Sub-cellular fractionation and ribosome association assays were done on single-round infected cells, as previously described for fractionation^6,37^ and ribosome association^10,22^. HIV-1 RNA splicing was assayed by Primer-ID tagged deep sequencing^10,38^. HIV-1 Gag protein expression levels were analyzed by Western blot. Western blot band intensity was quantified using Image J software. Released viral particles were quantified using an HIV-1 p24 antigen capture ELISA assay (Advanced Bioscience Laboratories #5421).

### RNA decay assays

The nascent RNA isolation method used is a combination of two protocols^23,24^. Single-round-infected cells were pulsed at 48 hpi with 150 μM 4SU for 1.5 hours, the cells then washed and resuspended in 4SU-free fresh RPMI. Cells were collected at 0, 2, and 4 h after 4SU wash out and RNA extracted using Trizol. 500 ng of MTSEA-biotin-XX (Biotium 89139-636) dissolved in 10 μl of Dimethyl formamide (Sigma D4551) was used to biotinylate 6 μg of RNA in a 50 μl reaction mixture with 20mM HEPES pH7.4 and 1mM EDTA at room temperature for 30 min. Excess biotin was removed by two rounds of chloroform extraction followed by isopropanol precipitation of RNA. 100 μg of streptavidin magnetic beads (NEB, S1420S) were pre-blocked with glycogen, then co-incubated with the Biotinylated-4SU+ RNA at room temperature for 15 min. The resulting RNA-bead complex was washed 3x with wash buffer (10mM Tris HCl pH7.4, 100mM NaCl, 1mM EDTA, 0.005% Tween-20). Elution was done twice with 25μl of freshly made elution buffer (20mM HEPES pH7.4, 100mM DTT, 1mM EDTA, 100mM NaCl, 0.05% Tween-20), followed by RNA purification with the Zymo RNA Clean & Concentrator-5 kit (#11-326). For the transcription stop method, single-round-infected cells were treated at 48 hpi with 5μg/ml Actinomycin D (Sigma A9415), and harvested 0, 2, 4, and 6 h later. For all RNA decay assays, viral RNA levels were assayed by qRT-PCR. Data were analyzed using the ddCt method, where readouts of viral RNA at time t (D1-A1 spliced RNA specific primer set) were normalized to β-Actin levels corrected to the expected RNA level prior to decay of t hours, utilizing the published β-Actin t_1/2_ in CEM cells of 13.5hrs (Ct of actin at time t is corrected by-t/13.5)^39^. Each β-actin-normalized viral RNA value was then calculated as the fold change from that detected at time point 0. Statistical analysis of the rate of RNA decay was done on the log2 transformation of each fold change value, using GraphPad Prism 8 software, comparing slopes of linear regression lines by analysis of covariance (ANCOVA).

## Supporting information

Supplemental Primer Table

## Data availability

All deep sequencing data have been deposited at the NCBI GEO database under accession number GSE142490.

## Code and reagent availability

The ΔNAT10 cell lines, all plasmid constructs, and data analysis Perl scripts are available upon request.

## Acknowledgements

We would like to thank Dr. Shalini Oberdoerffer for sharing the first batch of ac4C antibody and for valuable discussions, and Dr. Christopher Holley for advice and use of instruments. This research was funded in part by NIH grants R01-DA046111 to B.R.C. and U54-GM103297 to B.R.C. and R.S., along with a Duke University Center for AIDS Research (CFAR, P30-AI064518) pilot award to K.T. This research received infrastructure support from the Duke University CFAR, the UNC CFAR (P30-AI50410), and the UNC Lineberger Comprehensive Cancer Center (P30-CA16068). The following reagents were obtained through the NIH AIDS Reagent Program, Division of AIDS, NIAID, NIH: HIV-1 p24 Gag Monoclonal (#24-3) from Dr. Michael H. Malim and HIV-1 NL4-3 Infectious Molecular Clone (pNL4-3) from Dr. Malcolm Martin (# 114).

## Supplemental Material

**Fig. S1.**
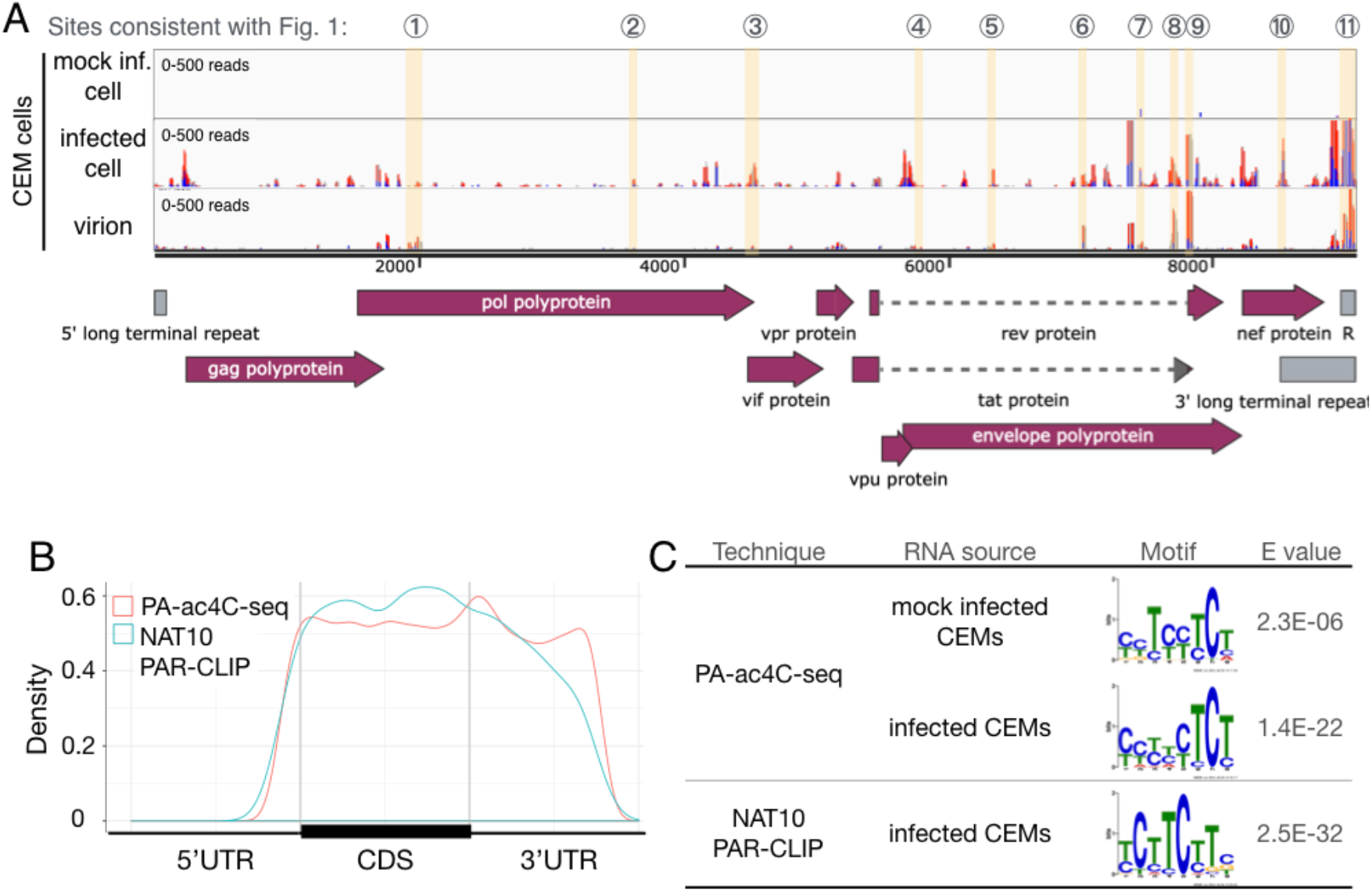
Distribution and sequence preference of cellular mRNA ac4C sites. (A) PA-ac4c-seq analysis of intracellular RNA and virion RNA recovered from CEM T cells (B) Metagene analysis of PA-ac4C-seq (from mock-infected CEM cells)-mapped ac4C sites and NAT10 PAR-CLIP-mapped NAT10-bound mRNA sites across the UTRs and coding regions (CDS) of cellular genes. (C) Enriched sequence motifs in ac4C (PA-ac4C-seq) and NAT10 (PAR-CLIP) binding sites. Data were analyzed on cDNA sequences, thus U residues in RNA are depicted as T residues.

**Fig S2.**
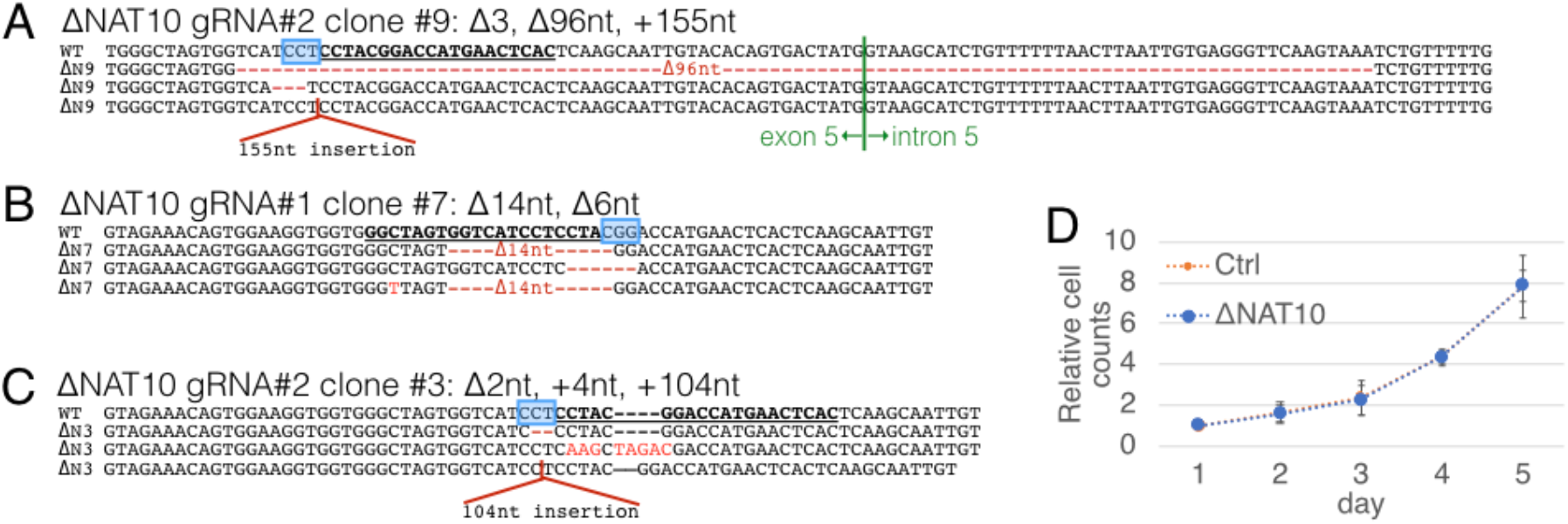
CRISPR-induced edits and viability of ΔNAT10 CEM cell lines. (A-C) The CRISPR-targeted genomic exon 5 region of three single cell clones of ΔNAT10 CEMs were cDNA cloned and subjected to Sanger sequencing. The mutated sequences of ΔNAT10 clones #9, #7, and #3 shown here. The Cas9 PAM, 5’-NGG-3’ isshown in blue boxes, guide RNA targeted sequence in underlined bold text, indels and substitutions shown in red. Unless specifically stated, all assays used ΔNAT10 clone #9. (D) Ctrl and ΔNAT10 CEMs were cultured side by side and the cells counted for 5 days. n=3, error bar=SD.

**Fig S3.**
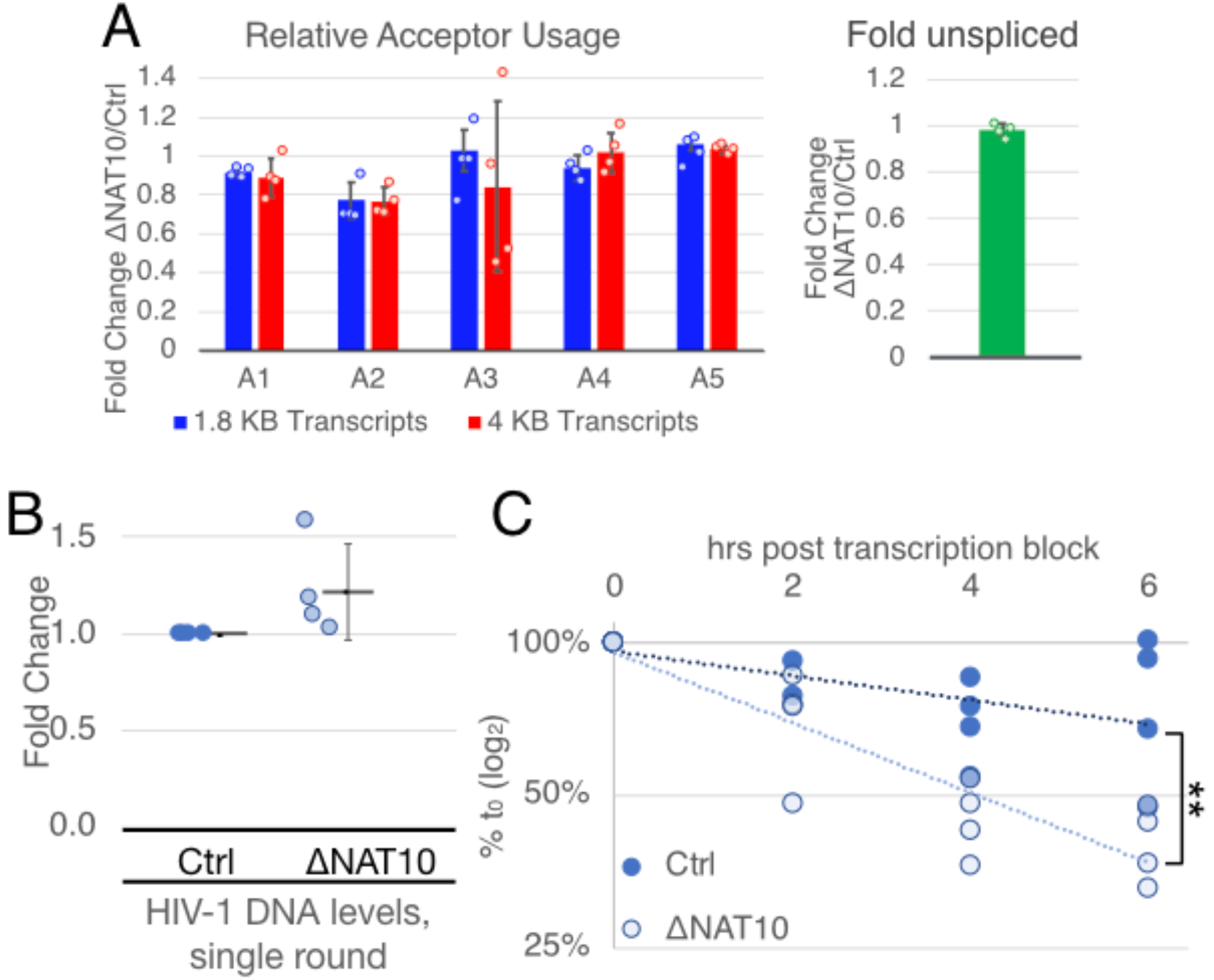
Splicing, viral DNA levels, and RNA stability in single-round infected Ctrl & ΔNAT10 CEM cells. (A-B) Viral transcript splice forms quantified by Primer-ID tagged deep sequencing on incompletely spliced (~4kb), completely spiced (~1.8kb), or unspliced transcripts, plotted as fold change of splice acceptor usage (A1-5) of ΔNAT10 over Ctrl, n=4. (B) Viral DNA levels in single-round infected Ctrl and ΔNAT10 CEMs, 48 hpi, quantified by qPCR using an LTR U3 primer set, n=4. (C) Viral RNA stability assayed in single-round infected Ctrl and ΔNAT10 CEMs by blocking transcription with Actinomycin D (ActD) at 2 dpi and assaying the viral RNA levels at the shown time points after ActD treatment by qRT-PCR. Slopes of regression lines compared by ANCOVA, n=4, **p=0.0025.

**Fig S4.**
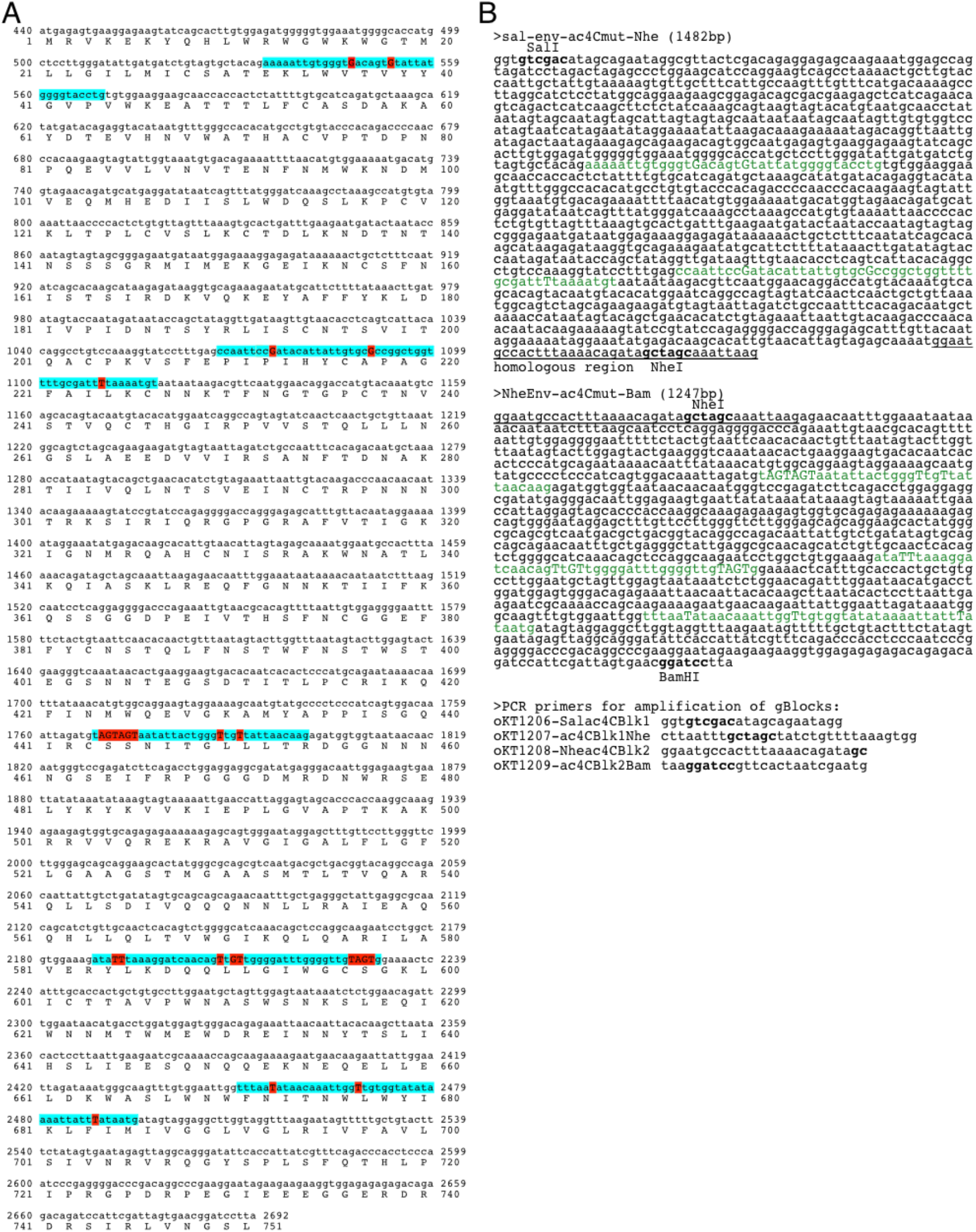
Sequence and construction of ac4C silent point mutant virus. (A) Coding sequence and translated amino acid sequence of the mutated env gene. Mapped ac4C sites highlighted in blue, mutated nucleotides capitalized in red. (B) gBlocks and amplification primers used to construct the mutant virus. Mapped ac4C sites in green, mutated nucleotides capitalized. A region of homology was designed between the two gBlocks (underlined) to facilitate recombinant PCR. Restriction enzyme sites used for cloning depicted in bold.

## References

1. Roundtree, I.A., Evans, M.E., Pan, T. & He, C. Dynamic RNA modifications in gene expression regulation. Cell 169, 1187–1200 (2017).

2. Kennedy, E.M., Courtney, D.G., Tsai, K. & Cullen, B.R. Viral epitranscriptomics. J Virol 91, e02263–16 (2017).

3. Kennedy, E.M. et al. Posttranscriptional m(6)A editing of HIV-1 mRNAs enhances viral gene expression. Cell Host Microbe 19, 675–85 (2016).

4. Lichinchi, G. et al. Dynamics of the human and viral m6A RNA methylomes during HIV-1 infection of T cells. Nature Microbiology 1, 16011 (2016).

5. Courtney, D.G. et al. Epitranscriptomic enhancement of influenza A virus gene expression and replication. Cell Host Microbe 22, 377–386 e5 (2017).

6. Tsai, K., Courtney, D.G. & Cullen, B.R. Addition of m6A to SV40 late mRNAs enhances viral structural gene expression and replication. PLoS Pathog 14, e1006919 (2018).

7. Xue, M. et al. Viral N(6)-methyladenosine upregulates replication and pathogenesis of human respiratory syncytial virus. Nat Commun 10, 4595 (2019).

8. Hao, H. et al. N6-methyladenosine modification and METTL3 modulate enterovirus 71 replication. Nucleic Acids Res 47, 362–374 (2019).

9. Ye, F., Chen, E.R. & Nilsen, T.W. Kaposi’s Sarcoma-Associated Herpesvirus Utilizes and Manipulates RNA N6-Adenosine Methylation To Promote Lytic Replication. J Virol 91(2017).

10. Courtney, D.G. et al. Epitranscriptomic Addition of m(5)C to HIV-1 Transcripts Regulates Viral Gene Expression. Cell Host Microbe 26, 217–227 e6 (2019).

11. Hesser, C.R., Karijolich, J., Dominissini, D., He, C. & Glaunsinger, B.A. N6-methyladenosine modification and the YTHDF2 reader protein play cell type specific roles in lytic viral gene expression during Kaposi’s sarcoma-associated herpesvirus infection. PLoS Pathog 14, e1006995 (2018).

12. Arango, D. et al. Acetylation of Cytidine in mRNA Promotes Translation Efficiency. Cell 175, 1872–1886 e24 (2018).

13. Balmus, G. et al. Targeting of NAT10 enhances healthspan in a mouse model of human accelerated aging syndrome. Nat Commun 9, 1700 (2018).

14. Larrieu, D., Britton, S., Demir, M., Rodriguez, R. & Jackson, S.P. Chemical inhibition of NAT10 corrects defects of laminopathic cells. Science 344, 527–32 (2014).

15. Ringeard, M., Marchand, V., Decroly, E., Motorin, Y. & Bennasser, Y. FTSJ3 is an RNA 2’-O-methyltransferase recruited by HIV to avoid innate immune sensing. Nature 565, 500–504 (2019).

16. McIntyre, W. et al. Positive-sense RNA viruses reveal the complexity and dynamics of the cellular and viral epitranscriptomes during infection. Nucleic Acids Res 46, 5776–5791 (2018).

17. Sinclair, W.R. et al. Profiling Cytidine Acetylation with Specific Affinity and Reactivity. ACS Chem Biol 12, 2922–2926 (2017).

18. Ito, S. et al. Human NAT10 is an ATP-dependent RNA acetyltransferase responsible for N4-acetylcytidine formation in 18 S ribosomal RNA (rRNA). J Biol Chem 289, 35724–30 (2014).

19. Sharma, S. et al. Yeast Kre33 and human NAT10 are conserved 18S rRNA cytosine acetyltransferases that modify tRNAs assisted by the adaptor Tan1/THUMPD1. Nucleic Acids Res 43, 2242–58 (2015).

20. Hafner, M. et al. Transcriptome-wide identification of RNA-binding protein and microRNA target sites by PAR-CLIP. Cell 141, 129–41 (2010).

21. Mefferd, A.L., Bogerd, H.P., Irwan, I.D. & Cullen, B.R. Insights into the mechanisms underlying the inactivation of HIV-1 proviruses by CRISPR/Cas. Virology 520, 116–126 (2018).

22. Subtelny, A.O., Eichhorn, S.W., Chen, G.R., Sive, H. & Bartel, D.P. Poly(A)-tail profiling reveals an embryonic switch in translational control. Nature 508, 66–71 (2014).

23. Dolken, L. et al. High-resolution gene expression profiling for simultaneous kinetic parameter analysis of RNA synthesis and decay. RNA 14, 1959–72 (2008).

24. Duffy, E.E. et al. Tracking Distinct RNA Populations Using Efficient and Reversible Covalent Chemistry. Mol Cell 59, 858–66 (2015).

25. Sanjana, N.E., Shalem, O. & Zhang, F. Improved vectors and genome-wide libraries for CRISPR screening. Nat Methods 11, 783–4 (2014).

26. Simon, J.H. et al. The Vif and Gag proteins of human immunodeficiency virus type 1 colocalize in infected human T cells. J Virol 71, 5259–67 (1997).

27. Adachi, A. et al. Production of acquired immunodeficiency syndrome-associated retrovirus in human and nonhuman cells transfected with an infectious molecular clone. J Virol 59, 284–91 (1986).

28. Chen, K. et al. High-resolution N(6)-methyladenosine (m(6) A) map using photo-crosslinking-assisted m(6) A sequencing. Angew Chem Int Ed Engl 54, 1587–90 (2015).

29. Langmead, B. Aligning short sequencing reads with Bowtie. Curr Protoc Bioinformatics **Chapter 11**, Unit 11 7 (2010).

30. Li, H. et al. The Sequence Alignment/Map format and SAMtools. Bioinformatics 25, 2078–9 (2009).

31. Robinson, J.T. et al. Integrative genomics viewer. Nat Biotechnol 29, 24–6 (2011).

32. Liu, J. et al. A METTL3-METTL14 complex mediates mammalian nuclear RNA N6-adenosine methylation. Nat Chem Biol 10, 93–5 (2014).

33. Corcoran, D.L. et al. PARalyzer: definition of RNA binding sites from PAR-CLIP short-read sequence data. Genome Biol 12, R79 (2011).

34. Dominissini, D., Moshitch-Moshkovitz, S., Salmon-Divon, M., Amariglio, N. & Rechavi, G. Transcriptome-wide mapping of N(6)-methyladenosine by m(6)A-seq based on immunocapturing and massively parallel sequencing. Nat Protoc 8, 176–89 (2013).

35. Olarerin-George, A.O. & Jaffrey, S.R. MetaPlotR: a Perl/R pipeline for plotting metagenes of nucleotide modifications and other transcriptomic sites. Bioinformatics 33, 1563–1564 (2017).

36. Bieniasz, P.D., Fridell, R.A., Anthony, K. & Cullen, B.R. Murine CXCR-4 is a functional coreceptor for T-cell-tropic and dual-tropic strains of human immunodeficiency virus type 1. J Virol 71, 7097–100 (1997).

37. Malim, M.H., Hauber, J., Le, S.Y., Maizel, J.V. & Cullen, B.R. The HIV-1 rev trans-activator acts through a structured target sequence to activate nuclear export of unspliced viral mRNA. Nature 338, 254–7 (1989).

38. Emery, A., Zhou, S., Pollom, E. & Swanstrom, R. Characterizing HIV-1 Splicing by Using Next-Generation Sequencing. J Virol 91, e02515–16 (2017).

39. Leclerc, G.J., Leclerc, G.M. & Barredo, J.C. Real-time RT-PCR analysis of mRNA decay: half-life of Beta-actin mRNA in human leukemia CCRF-CEM and Nalm-6 cell lines. Cancer Cell Int 2, 1 (2002).

